# Does non-invasive vagus nerve stimulation affect heart rate variability? A living and interactive Bayesian meta-analysis

**DOI:** 10.1101/2021.01.18.426704

**Authors:** Vinzent Wolf, Anne Kühnel, Vanessa Teckentrup, Julian Koenig, Nils B. Kroemer

## Abstract

Non-invasive brain stimulation techniques, such as transcutaneous auricular vagus nerve stimulation (taVNS), have considerable potential for clinical use. Beneficial effects of taVNS have been demonstrated on symptoms in patients with mental or neurological disorders as well as transdiagnostic dimensions, including mood and motivation. However, since taVNS research is still an emerging field, the underlying neurophysiological processes are not yet fully understood, and the replicability of findings on biomarkers of taVNS effects has been questioned. Here, we perform a living Bayesian random effects meta-analysis to synthesize the current evidence concerning the effects of taVNS on heart rate variability (HRV), a candidate biomarker that has, so far, received most attention in the field. To keep the synthesis of evidence transparent and up to date as new studies are being published, we developed a Shiny web app that regularly incorporates new results and enables users to modify study selection criteria to evaluate the robustness of the inference across potential confounds. Our analysis focuses on 17 single-blind studies comparing taVNS versus sham in healthy participants. These newly synthesized results provide strong evidence for the null hypothesis (*g* = 0.011, *CI_shortest_* = [−0.103, 0.125], *BF_01_* = 25.587), indicating that acute taVNS does not alter HRV compared to sham. To conclude, based on a synthesis of the available evidence to date, there is no support for the hypothesis that HRV is a robust biomarker for acute taVNS. By increasing transparency and timeliness, we believe that the concept of living meta-analyses can lead to transformational benefits in emerging fields such as non-invasive brain stimulation.

## Introduction

Emerging fields such as non-invasive brain stimulation provide unique challenges in synthesizing evidence. First described two decades ago (Ventureyra, 2000), transcutaneous auricular vagus nerve stimulation (taVNS) is a novel, cost-effective alternative to invasive cervical vagus nerve stimulation (iVNS), which is an FDA-approved treatment for epilepsy, treatment-resistant depression and morbid obesity (Badran et al., 2018). In contrast to iVNS, taVNS is performed with electrodes placed at stimulation sites located at the ear. Crucially, the application of taVNS has shown similar beneficial results compared to iVNS, for example in the reduction of symptoms in patients with depression (Fang et al., 2016) and altered early visual processing of negative emotional stimuli in adolescent depression (Koenig et al., 2019). Likewise, beneficial effects of taVNS were also found for chronic pain (Napadow et al., 2012) and epilepsy (Aihua et al., 2014). These similarities in effects might be explained by the similarity of brain network activation achieved by iVNS and taVNS (Frangos et al., 2015; Kaniusas et al., 2019). Nonetheless, studies in healthy subjects also show diverse behavioral effects indicating multiple modes of taVNS action. For example, taVNS boosted mood after effort exertion, highlighting the vagus nerve’s role in modulating interoceptive feedback signals (Ferstl et al., 2021). In addition, taVNS modulated reinforcement learning (Kühnel et al., 2020), facilitated invigoration to work for food and monetary rewards (Neuser et al., 2020) and enhanced delay discounting for individuals with low positive mood (Steenbergen et al., 2020). Although taVNS has been shown to evoke a variety of intended behavioral effects in healthy as well as in clinical samples, the physiological processes underlying the effects of taVNS are still largely elusive (Burger et al., 2020). Lack of convergence across behavioral studies and multiple theories on the mode of action make it imperative to identify a reliable biomarker for successful vagal activation via taVNS.

A reliable biomarker for successful vagus nerve activation would be crucial as this could provide “information on the extent, to which the vagus nerve is actually being stimulated” (Burger et al., 2020). Especially in clinical settings, a biomarker is urgently needed to distinguish between responders and non-responders to taVNS interventions (Yap et al., 2020). Although multiple potential biomarkers have been proposed, most research has focused on heart rate variability (HRV). As major nerve of the parasympathetic system, the vagus nerve innervates the heart among other organs (Kaniusas et al., 2019; Schachter & Saper, 1998). Both the left and right vagus innervate the heart’s sinoatrial node, which serves as the heart’s natural pacemaker (Monfredi et al., 2010). Thereby, taVNS may affect heart rate and HRV via vagal efferents. Although specific indices of HRV have been shown to reliably index parasympathetic vagal modulation, heart rate is not only influenced by vagal activity. Since heart rate incorporates sympathetic and parasympathetic activity, it should not be interpreted as a valid marker of vagal activity. Crucially, the right vagus innervates the sinoatrial node more extensively (Ardell & Randall, 1986; Guiraud et al., 2016; Kaniusas et al., 2019) which is consistent with anatomical findings that invasive stimulation of the right vagus was found to increase HRV in animals (Lee et al., 2018; Yoshida et al., 2018), whereas left-sided stimulation produced mixed effects (Martlé et al., 2014; Samniang et al., 2016). In humans, usually only the left side is stimulated due to concerns about cardiac side effects (e.g. Borges et al., 2019; Burger et al., 2019a; Burger et al., 2016). However, the strength of potential side-specific effects as observed in animal models is currently under debate. For instance, De Couck and colleagues compared different stimulation protocols and found an increase in HRV only for right-sided stimulation (De Couck et al., 2017). In contrast, other authors argue against practically relevant differences between stimulation sides, as the vagal nerve endings in the ear are almost exclusively afferent (Kaniusas et al., 2019). As a result, information from both sides might be integrated in the brainstem centrally, before influencing cardiac activity efferently (Chen et al., 2015). Taken together, given the vital role of the vagus nerve within the parasympathetic system, an increase in HRV is an anatomically plausible—albeit debated—candidate biomarker.

Despite the anatomical plausibility of HRV as a biomarker for taVNS, reported acute effects of taVNS on HRV are mixed. For example, some studies found a significant decrease in the low frequency to high frequency (LF/HF) ratio in active taVNS compared to sham, indicating an increase in HRV, but no differences in high-frequency (HF) HRV (Antonino et al., 2017; Clancy et al., 2014). One study with patients showing diastolic dysfunction found increased HRV after taVNS compared to sham for both HF-HRV and LF/HF ratio (Tran et al., 2019). Lamb and colleagues found an increase in HRV indexed by respiratory sinus arrhythmia (RSA) in healthy controls and patients with either posttraumatic stress disorder (PTSD) or PTSD and mild traumatic brain injury (Lamb et al., 2017). However, many studies examining healthy participants did not find an increase in HRV (e.g. Burger et al., 2019a, 2019b, Borges et al., 2019; Teckentrup et al., 2020). Evidence from narrative reviews is inconclusive as well. Kaniusas and colleagues (Kaniusas et al., 2019) argued for the presence of an effect of taVNS on HRV, de-emphasizing studies that did not find such effects. Burger and colleagues (Burger et al., 2020) reviewed different candidate biomarkers for taVNS, but they did not draw a definitive conclusion about HRV. Likewise, Butt and colleagues (Butt et al., 2020) focused on the anatomical basis of tVNS and did not find conclusive support for a modulation of HRV. There are a number of possible explanations for the heterogeneity of effects. First, taVNS primarily excites Aβ-fibers, which do not innervate the heart (De Couck et al., 2017; Safi et al., 2016; Vuckovic et al., 2008). Second, even effects of iVNS on HRV show greater inconsistency in human compared to animal studies (Burger et al., 2020). Third, the auricular branch of the vagus nerve projects afferently to the nucleus of the solitary tract (NTS), which influences heart rate only indirectly via the nucleus ambiguous and the dorsal vagal nucleus (Murray et al., 2016). Fourth, large methodological differences between studies such as different stimulation devices, stimulations sides and sites, experimental designs, reported HRV parameters and stimulation protocols reduce comparability between studies. One striking example is the use of different control conditions. While most studies compare active taVNS versus earlobe sham as independent variable as recommended (Farmer et al., 2020), some studies compare against a non-stimulation control condition (Tobaldini et al., 2019) or a non-stimulation sham condition, where the electrode is placed on the ear, but no electrical current is applied (De Couck et al., 2017; Sclocco et al., 2020). Consequently, to evaluate convergence across a set of emerging evidence from heterogeneous studies, we need improved meta-analytic tools to provide dependable results for future research and applications in clinical use settings.

Conventionally, reviews (narrative or systematic) and meta-analyses are used to provide a definitive answer to a pressing question. However, these conventional formats have been criticized because readers are “at the mercy of the author’s discretion” (Ahern et al., 2020). Additionally, authors are often subject to the reviewer bias (Ernst, 1992), thus being unable to detach themselves from their previous experiences and assumptions. To tackle these issues, we developed a Shiny web application (Chang et al., 2019) accompanying our meta-analysis, based on the example of Ahern and colleagues (Ahern et al., 2020). To substantially improve transparency and robustness of the inference, users can select inclusion criteria as well as prior specifications for both effect and heterogeneity based on their perspective and evaluate whether the data provides a conclusive result or not in light of their assumptions. Moreover, this meta-analysis format allows for future research to be added as it emerges, keeping the results up to date. The application can be opened via the link: https://vinzentwolf.shinyapps.io/taVNSHRVmeta/. Therefore, our living meta-analysis helps to clarify if taVNS acutely increases HRV compared to sham while adapting to the growing body of literature in an emerging field and therefore adds important insight to the research on neurophysiological effects of taVNS.

## Methods

### HRV measurement

HRV is a commonly used index for cardiac vagal activity, reflecting parasympathetic contributions to cardiac regulation (Hayano et al., 1991; Laborde et al., 2017). However, multiple indices are used to index HRV (Burger et al., 2020; Electrophysiology, 1996) and were therefore included in the present analysis as dependent variables. The standard deviation of all R-to-R intervals (SDNN) reflects cyclic components responsible for heart rate variability. The root mean square of successive differences (RMSSD), the percentage of successive normal sinus RR intervals more than 50ms apart (pNN50) and SDNN represent HRV in the time-domain. Critically, SDNN is thought to reflect both, sympathetic and parasympathetic influences, whereas RMSSD and pNN50 are thought to represent vagally-mediated HRV. Further, SDNN depends on the length of the recording period, thus making it difficult to compare SDNN measures obtained from recordings of different durations (Electrophysiology, 1996). High frequency power (HF, >0.15Hz) indicates vagal activity and thus reflects HRV in the frequency domain (Laborde et al., 2017). Additionally, the ratio of low to high frequencies (LF/HF) is included in the analysis, although it reflects a mix of sympathetic and vagal activity and its physiological origin is disputed (Heathers & Goodwin, 2017). Nonetheless, many researchers are still reporting the LF/HF-ratio as dependent variable (e.g. Bretherton et al., 2019; Clancy et al., 2014; De Couck et al., 2017; Tobaldini et al., 2019; Tran et al., 2019; Villani et al., 2019). Another measure assessing HRV included in our meta-analysis is cardiac vagal tone (CVT) calculated using a process called phase-shift demodulation (Frøkjaer et al., 2016; Juel et al., 2017). Lastly, one study measured respiratory sinus arrhythmia (RSA) as an indicator of HRV (Lamb et al., 2017) and was included, too.

### Inclusion and exclusion criteria – Shiny app

To be included in the Shiny app, studies had to measure at least one of the following HRV parameters: RMSSD, SDNN, pNN50, HF-HRV, LF/HF-Ratio, CVT, or RSA acutely during stimulation of the tragus, cymba conchae or cavum conchae. Also, they needed to compare an active taVNS condition against a sham condition or a pre-stimulation baseline. All stimulation sides (left, right, bilaterally) were included. Studies examining healthy and clinical participants were eligible for inclusion. Since research on taVNS started only two decades ago, we did not apply any restrictions regarding the time period in which studies needed to be published. Further, we did not make any geographical or cultural restrictions nor any restrictions regarding the participants’ demographics.

### Inclusion and exclusion criteria – exemplary analysis

For the analysis presented in this paper, we restricted the sample to only include studies with a high-quality design. Therefore, only at least single-blind studies comparing active taVNS against sham were included, but not pre-stimulation baseline comparisons. SDNN and LF/HF ratio were excluded as HRV parameters, as they do not index vagal activity. Therefore, included studies had to measure at least one of the following parameters: RMSSD, pNN50, HF-HRV, CVT, or RSA. To account for possible confounds because of clinical disorders influencing HRV (Farmer et al., 2020; Kemp et al., 2010; Thayer et al., 2010), we additionally excluded studies examining patient samples from the analyses presented here.

### Search strategies

We performed database searches in Web of Science, Science Direct, PsycInfo, PubMed and Google Scholar on 23/07/2020 with the following keywords: *“(HRV OR heart rate variability OR parasympathetic OR root means square of successive differences OR rmssd OR vagal tone OR heart rate OR hr OR pNN50) AND (transcutaneous auricular vagus nerve stimulation OR transcutaneous vagus nerve stimulation OR tVNS OR taVNS)”*. Additionally, we screened three reviews (Burger et al., 2020; Farmer et al., 2020; Redgrave et al., 2018) to identify eligible reports. Lastly, we sent e-mails to the members of the transcutaneous vagus nerve stimulation consensus group (Farmer et al., 2020), asking for unpublished HRV data from taVNS experiments. If a paper indicated that HRV was indeed measured but the results were reported insufficiently or the raw data was not published, we contacted the authors via e-mail. Records were screened and assessed for eligibility following the recommendations of the PRISMA group (Moher et al., 2009). No reports in languages other than English were included.

### Coding procedures

Reports were coded by two coders and differences were clarified through discourse. We coded design type (within-subjects, between-subjects), blindness (double blind, single blind, transparent) and control type (sham, baseline) regarding the assessment of study quality. If necessary, missing values were calculated. Two studies reported means and standard errors (SE) only (Bretherton et al., 2019; Clancy et al., 2014), hence we calculated standard deviations (SD) with *SD = SE*sqrt(N_group_)*. One study (Tobaldini et al., 2019) reported medians and inter quartile ranges (IQR), hence means were estimated with *mean = (IQR_lower_ + median + IQR_upper_)/3* and SDs with *SD = (IQR_upper_ – IQR_lower_)/1.206*, with *1.206* being a constant depending on the sample size (Wan et al., 2014). Further, for two papers (Burger et al., 2019a; Burger et al., 2019b), we calculated effect sizes from the results of the corresponding independent t-tests using the *esc* package (Lüdecke, 2019) in R 4.0.2 (R Core Team, 2020) using RStudio (RStudio Team, 2020).

### Design of the Shiny app

The full analysis including the inclusion criteria can be found in the accompanying Shiny app as well. When initially opening the app, the default inclusion criteria correspond to the inclusion criteria reported in this paper. Since results might still depend on different HRV and stimulation parameter choices as well as on inclusion/exclusion criteria and prior settings, the accompanying Shiny app provides the possibility to perform the analysis according to the user’s preferences. The sidebar allows for individual configuration of the selection criteria. Inclusion criteria can be modified according to the design, type of control condition, calculated HRV parameters, sample, gender, age, blindness of the studies, and publication year. Moreover, the taVNS specifications panel allows choices about stimulation side, stimulation site, stimulation duration, intensity and stimulation device used. Pressing *“Re-Calculate Meta-Analysis”* will rerun the analysis with the newly set inclusion criteria. In the app’s *“Study overview”* panel, included studies, their effect size and relevant study parameters are briefly summarized. If less than four studies are included due to too strict inclusion criteria, a warning message gets displayed. The plots depicted in the *“Outlier check”*, *“Forest plot”* and *“Funnel plot”* panels have the same properties as their respective counterparts above. If a proper τ prior is chosen, Bayes factors for heterogeneity and effect will be displayed in the *“Statistics”* panel. The marginal posterior summary provides descriptive statistics and 95% shortest credible intervals for the posterior distributions of heterogeneity, taVNS effect and prediction. Joint and marginal maximum likelihood (ML) and maximum a posteriori (MAP) estimates are displayed in the *“Statistics”* panel as well. *“Full texts screened”* provides a table showing all research articles which were assessed for eligibility for the meta-analysis. Users can request the addition of studies by submitting studies for inclusion. The *“Additional plots”* section shows the joint posterior density of heterogeneity and effect, distributions of effect prior, likelihood and posterior, and a distribution of the selected heterogeneity prior. Lastly, the optional *“Bayes factor robustness check”* provides a plot as in Figure 3a. R Code for the Shiny app can be found on GitHub^1^.

### Statistical methods

For each taVNS versus sham comparison, we calculated Hedges’ *g*s (Hedges & Olkin, 1985) as effect sizes based on means and SDs using the *metafor* package’s *escalc* function (Viechtbauer, 2010). Effect sizes were calculated for each taVNS versus pre-stimulation baseline comparison in the same way. These can be investigated in the Shiny app. Since most studies reported multiple HRV outcome measures, we aggregated effect sizes over different HRV parameters. By using this aggregation method, we were able to include more studies into one analysis, while other meta-analyses on HRV split the results for different HRV parameters (e.g. Kemp et al., 2010). Effect sizes were aggregated using the Borenstein-Hedges-Higgins-Rothstein (BHHR) method (Borenstein et al., 2009) as implemented in the *MAd* package (Hoyt & Del Re, 2015). This method takes within-study correlations into account and has shown to be the least biased and most precise method for aggregating within-study effect sizes (Del Re, 2015). Within-study correlation was set to the default setting of 0.5. Importantly, in the Shiny app, the filtering of studies based on the selected inclusion criteria is executed before the aggregation of within study effect sizes. For example, if a study reported RMSSD, pNN50 and HF-HRV as dependent measures, but the user decides to exclude pNN50, only RMSSD and HF-HRV effect sizes of this particular study will be aggregated. Consequently, this will result in differing effect size of each study depending on the selected inclusion criteria.

To test for the influence of acute taVNS on HRV across studies, we performed a Bayesian random-effects meta-analysis, using the *bayesmeta* package (Röver, 2020). The Bayesian approach is more suitable for meta-analyses including few studies, even as few as four (Harrer et al., 2019; Röver, 2020). Furthermore, in contrast to conventional frequentist results that allow only rejection of the null or alternative hypothesis, Bayesian approaches are able to find evidence for both null and alternative hypotheses. Bayesian results such as Bayes factors can be interpreted more intuitively than frequentist results, particularly if the results were to support the null hypothesis (Higgins et al., 2009; Röver, 2020). Additionally, Bayesian random-effect models allow for prediction intervals, enabling inferences about studies not included in the analysis (Higgins et al., 2009). We chose a random-effects model because it is implausible to assume fixed effects for studies varying in many aspects such as populations, HRV measures, and taVNS stimulation protocols. We report Bayes factors and shortest credible intervals (Liu et al., 2015) for both the effect of taVNS on HRV and the heterogeneity parameter. As Bayes factors depend on the priors defined for the effects, we calculated Bayes factors for different prior distributions with varying SDs, from “ultrawide” (the user selected SD +2), “wide” (the user selected SD +1), “user” (the user selected SD) to “narrow” (half the user selected SD), to evaluate the robustness of our results across narrow and wide priors. To assess potential publication bias, we used a funnel plot (Sterne et al., 2011). Since no systematic analyses regarding the effect of taVNS on HRV have been published so far, we used a weakly informative normally distributed prior with *mean = 0* and *SD = 1.5* for the effect of taVNS, μ. For the heterogeneity, τ, we selected a half Cauchy prior with a scale of 0.5, being a proper, informative prior (Röver, 2020) and thus recommended for routine use (Polson & Scott, 2012). However, the interactive analysis allows selecting different μ and τ priors, although changes to the prior settings should be performed with caution as they influence the results of the analysis (van Doorn et al., 2019). We used boxplot graphs to check for potential outliers that lie 1.5 IQRs or more above the 75% quantile or 1.5 IQRs or more below the 25% quantile, respectively.

## Results

Using a Bayesian random-effects meta-analysis approach, we analyzed if taVNS has an acute effect on HRV in healthy individuals compared to a sham control condition and provide a Shiny app to explore the analysis and results. We examined 182 papers for relevance after duplicates were removed. After screening of titles and abstracts, we assessed 70 full-text articles for eligibility. From these articles, we included 25 in the app. The most prominent reason for exclusion was the absence of HRV measurements (n = 30), as many studies only measured heart rate (e.g. Badran et al., 2018; Busch et al., 2013; Capone et al., 2015; Fischer et al., 2018; Yu et al., 2017). Another six studies measured only baseline HRV, but stopped measurement during active taVNS and were therefore excluded (e.g. Burger et al., 2016; Burger et al., 2017; Colzato et al., 2018). Additionally, four studies measured HRV, but reported the results insufficiently to derive estimates for the meta-analysis and thus were excluded. One study had neither a sham condition nor an HRV baseline measurement and was thus excluded (Paleczny et al., 2019). Furthermore, we excluded two studies which did not use taVNS but transcutaneous cervical vagus nerve stimulation (Brock et al., 2017) or transcutaneous electrical nerve stimulation on cervical acupuncture points (Chan et al., 2012), respectively. Lastly, two papers were excluded because they were study protocols and therefore reported no data (He et al., 2015; Li et al., 2015).

For the present analysis, we focused on high-quality designs in healthy participants. Therefore, we excluded 8 studies that are still part of the database provided by the Shiny app. Reasons for these exclusions were usage of SDNN and/or LF/HF ratio as only dependent measures (n = 1, Antonino et al., 2017), the use of clinical samples (n = 5) such as high worriers (Burger et al., 2019a), patients with hypertension (Sclocco et al., 2017), patients with diastolic dysfunction (Tran et al., 2019), patients with atrial fibrillation (Trobec et al., 2020) and patients with pancreatitis (Juel et al., 2017) or the absence of a sham control condition (n = 2) (Tobaldini et al., 2019; Weise et al., 2015). Therefore, 17 studies investigating the acute effects of taVNS on HRV compared to sham stimulation in healthy subjects are included in the following analysis. Across these studies, there was strong meta-analytic evidence for the null hypothesis (*g* = 0.011, *CI_shortest_* = [−0.103, 0.125], *BF_01_* = 25.587), suggesting no effect of taVNS on HRV (see Figure 1 and Table 1). Also, there was evidence for homogeneity in the reported effects (τ = 0.047, 95%CI = [0.000, 0.148], *BF_01_* = 9.152, Figure 2).

**Table 1:**
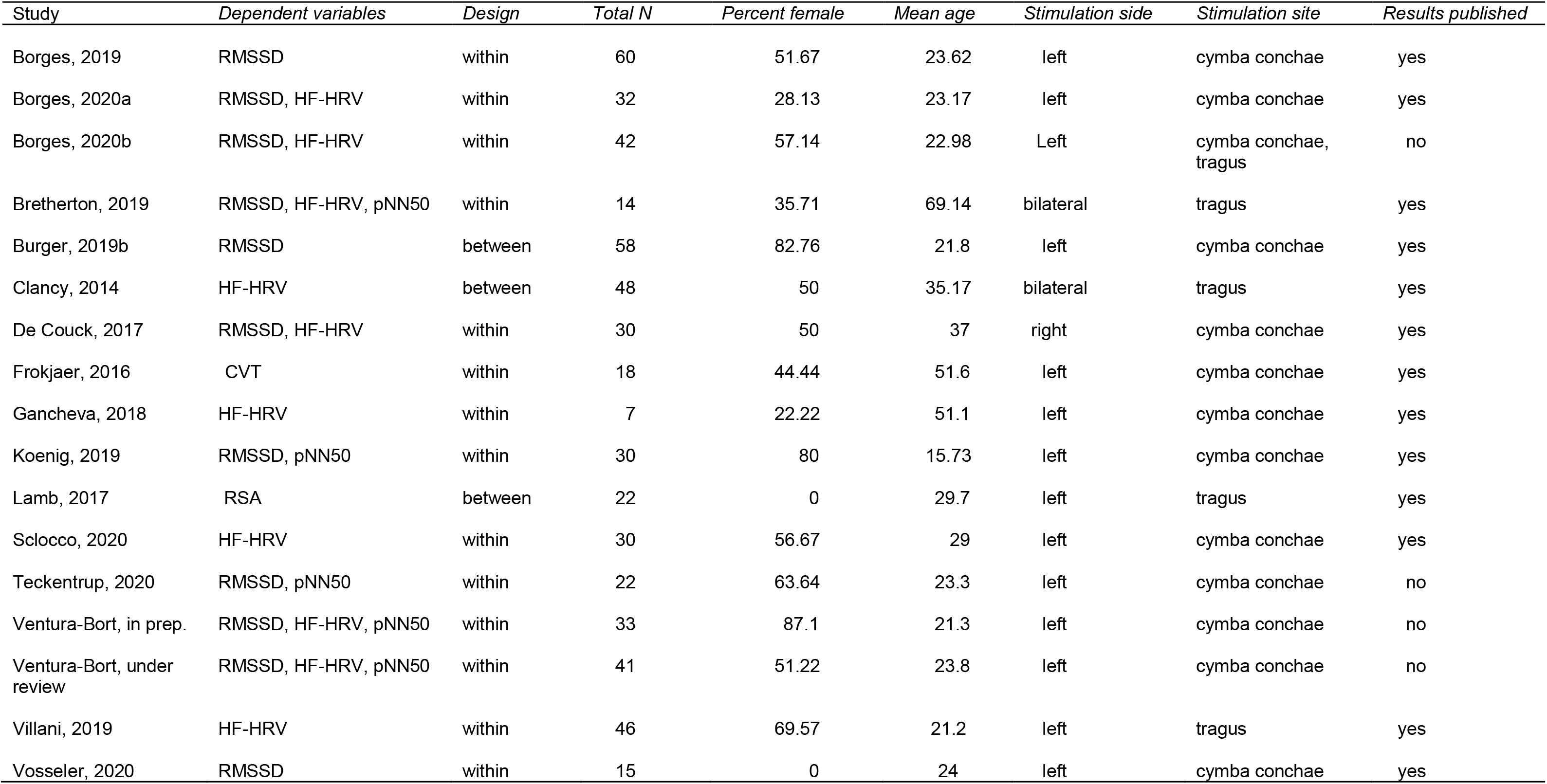
Studies included in the paper’s exemplary analysis. A more comprehensive table can be found in the Shiny app.

| Study | Dependent variables | Design | Total N | Percent female | Mean age | Stimulation side | Stimulation site | Results published |
| --- | --- | --- | --- | --- | --- | --- | --- | --- |
| Borges, 2019 | RMSSD | within | 60 | 51.67 | 23.62 | left | cymba conchae | yes |
| Borges, 2020a | RMSSD, HF-HRV | within | 32 | 28.13 | 23.17 | left | cymba conchae | yes |
| Borges, 2020b | RMSSD, HF-HRV | within | 42 | 57.14 | 22.98 | Left | cymba conchae, tragus | no |
| Bretherton, 2019 | RMSSD, HF-HRV, pNN50 | within | 14 | 35.71 | 69.14 | bilateral | tragus | yes |
| Burger, 2019b | RMSSD | between | 58 | 82.76 | 21.8 | left | cymba conchae | yes |
| Clancy, 2014 | HF-HRV | between | 48 | 50 | 35.17 | bilateral | tragus | yes |
| De Couck, 2017 | RMSSD, HF-HRV | within | 30 | 50 | 37 | right | cymba conchae | yes |
| Frokjaer, 2016 | CVT | within | 18 | 44.44 | 51.6 | left | cymba conchae | yes |
| Gancheva, 2018 | HF-HRV | within | 7 | 22.22 | 51.1 | left | cymba conchae | yes |
| Koenig, 2019 | RMSSD, pNN50 | within | 30 | 80 | 15.73 | left | cymba conchae | yes |
| Lamb, 2017 | RSA | between | 22 | 0 | 29.7 | left | tragus | yes |
| Sclocco, 2020 | HF-HRV | within | 30 | 56.67 | 29 | left | cymba conchae | yes |
| Teckentrup, 2020 | RMSSD, pNN50 | within | 22 | 63.64 | 23.3 | left | cymba conchae | no |
| Ventura-Bort, in prep. | RMSSD, HF-HRV, pNN50 | within | 33 | 87.1 | 21.3 | left | cymba conchae | no |
| Ventura-Bort, under review | RMSSD, HF-HRV, pNN50 | within | 41 | 51.22 | 23.8 | left | cymba conchae | no |
| Villani, 2019 | HF-HRV | within | 46 | 69.57 | 21.2 | left | tragus | yes |
| Vosseler, 2020 | RMSSD | within | 15 | 0 | 24 | left | cymba conchae | yes |

**Figure 1:**
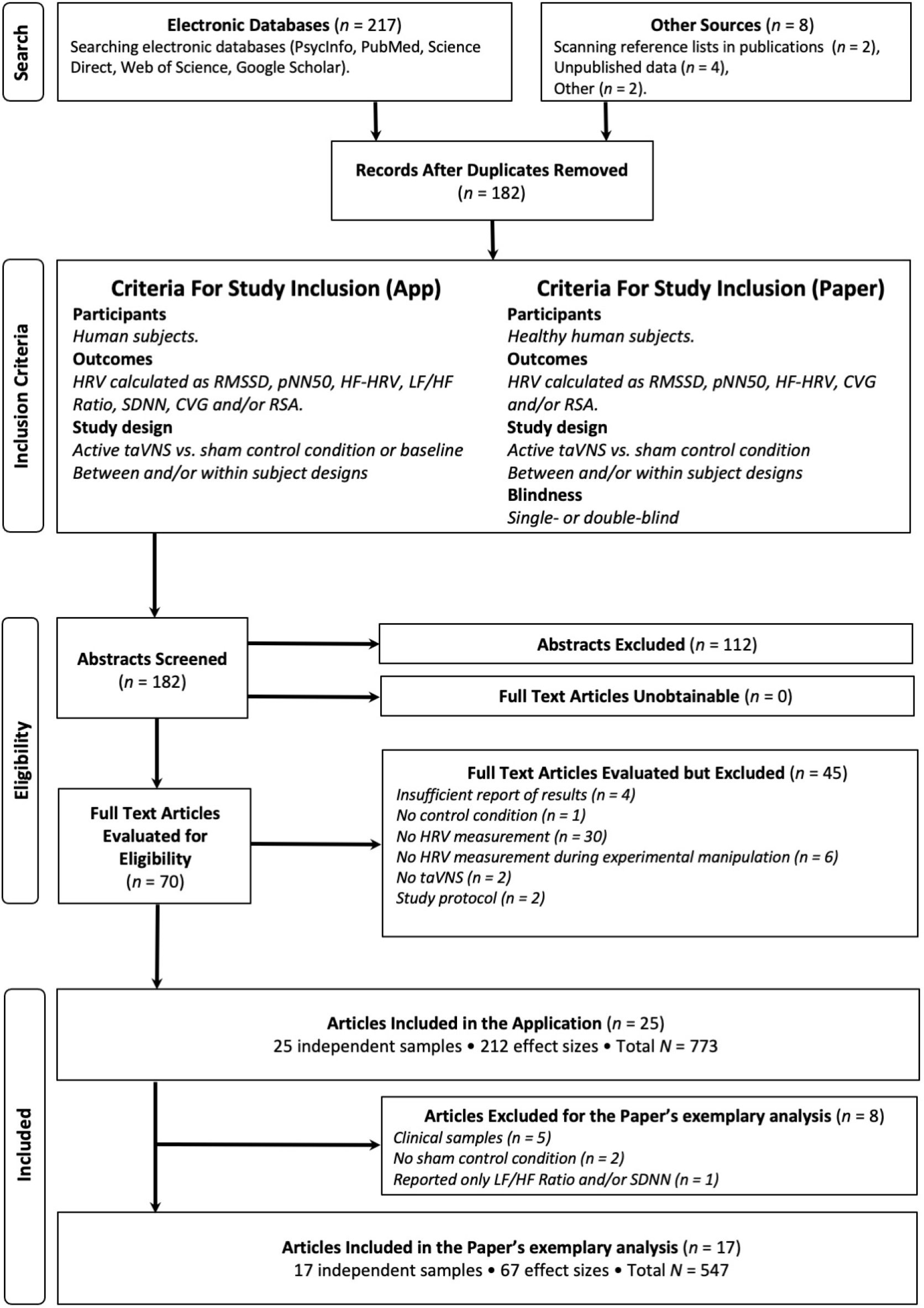
Exclusion process for articles included in the application and articles included in the paper’s exemplary analysis, following PRISMA recommendations (Moher et al., 2009).

**Figure 2:**
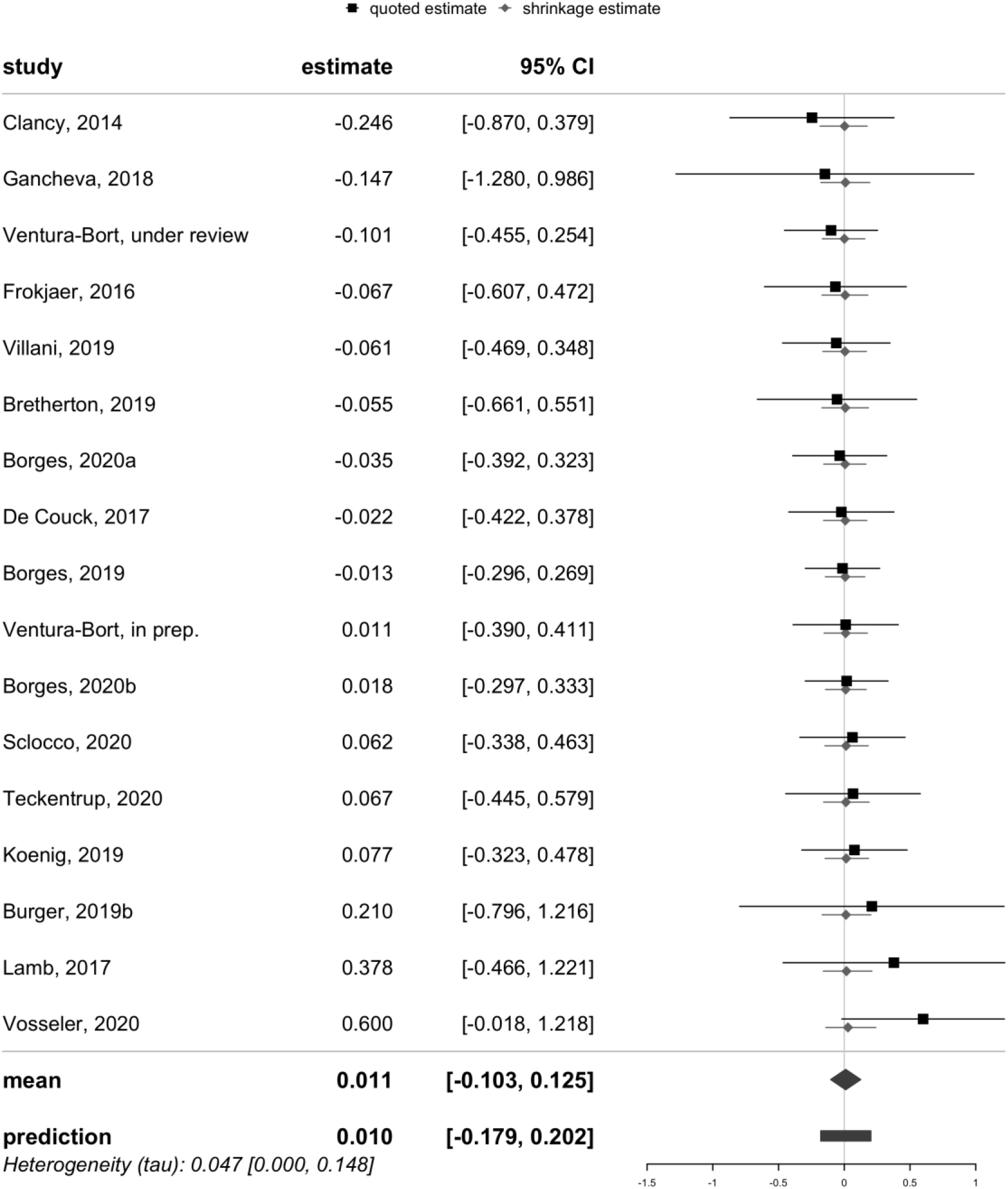
Forest plot of all included articles, their effect size estimates and 95% shortest credible intervals. Additionally, shrinkage of the individual studies’ posterior estimates towards the mean as a function of the heterogeneity are displayed (Röver, 2020).

Crucially, robustness analysis of the results across different prior widths showed at least substantial evidence for the null hypothesis (Figure 3), suggesting that the absence of an effect of taVNS was not dependent on prior settings. In line with the prior robustness analysis, we observed congruent results using a conventional frequentist analysis (*b* = 0.009, *SE* = 0.055, 95%CI = [−0.098, 0.116], *p* = ․865). Two studies were flagged as potential outliers (Lamb et al., 2017; Vosseler et al., 2020) with effect sizes of *g* = 0.378 and *g* = 0.6 (Figure 3b), respectively. However, excluding them did not alter the results (*g* = −0.016, 95%CI = [−0.132, 0.101], *BF_01_* = 24.632). The funnel plot (Figure 3c) indicates little evidence of publication bias.

**Figure 3:**
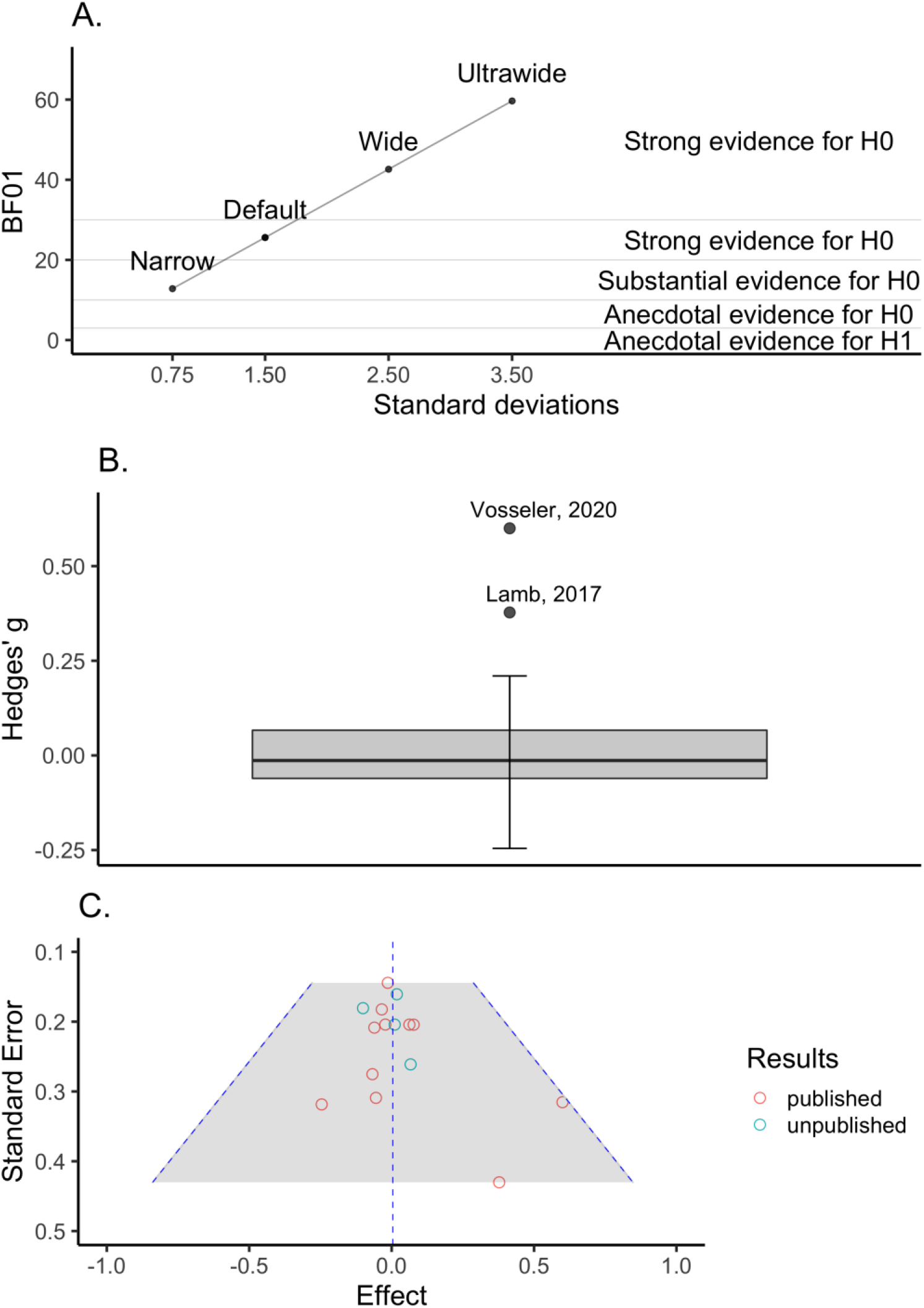
Converging evidence across 17 studies suggests that there is no effect of transcutaneous auricular vagus nerve stimulation (taVNS) on heart rate variability (HRV). a) Bayes Factor robustness plot. Bayes Factors are plotted across prior distributions differing in width. b) Boxplot graph identifying two studies (Lamb, 2017; Vosseler, 2020) as outliers. c) Funnel plot indicating little evidence of publication bias. Different colors of the points indicate whether the results were published or not.

## Discussion

The vagus nerve is an important part of the parasympathetic nervous system that can be stimulated non-invasively via the auricular branch at the ear. To measure hypothesized changes in vagal activity, HRV has been repeatedly suggested to serve as useful biomarker, but a systematic synthesis of the existing evidence is needed to support such assumptions. Here, using an interactive, living meta-analysis, we demonstrated a powerful way of synthesizing evidence from newly emerging research fields. The Bayesian random-effects meta-analysis provided strong evidence for the absence of an effect of taVNS compared to sham on HRV. Since the effect was absent across a wide range of specifications, our results indicate that HRV may not be a suitable biomarker or positive control outcome of effective taVNS – based on the existing studies in the field. Notwithstanding, additional research may identify more narrow conditions where taVNS could elicit a robust effect on HRV. Since these studies can be seamlessly integrated in the Shiny app, the analysis will gain depth and precision as the research field of taVNS will continue to grow. More broadly, our analysis may serve as a worked example for emerging fields in general as there are many other candidate biomarkers that would benefit from a principled and systematic analysis to quickly inform research on their suitability, for example, for clinical studies or patient monitoring.

Our results suggest that taVNS does not influence HRV compared to sham stimulation. In contrast to narrative reviews that were unable to draw clear conclusions (Burger et al., 2020; Butt et al., 2020) or argued for the existence of a taVNS effect on HRV (Kaniusas et al., 2019), we only included studies in healthy participants with a sufficient quality of the design in the default analysis such as sham control and (single-)blinded designs. Importantly, the derived evidence suggested homogeneity across studies and little indication of publication bias. This homogeneity and lack of publication bias is plausible because HRV was often not the primary outcome of the reported study, further strengthening the generalizability of our findings. Large methodological differences in stimulation duration, stimulation sides, sites and protocol between studies might attenuate the average effect-size estimate by introducing design-related noise. However, in light of the homogeneity analysis, such an explanation for the absence of a taVNS effect on HRV seems unlikely. Thus, according to the currently available data on taVNS-induced modulation of HRV, there is conclusive support for the absence of an effect across a wide range of inclusion settings and outcome measures. Notably, additional confounds on the subject and study level may be added in the future as more data become available. Thus, although the present findings are not in favor of HRV as biomarker for taVNS, future research including HRV measures may provide more nuanced insight.

Arguably, there are neurobiologically plausible explanations for the lack of robust taVNS effects on HRV. taVNS primarily excites Aβ-fibers, which do not innervate the heart (De Couck et al., 2017; Vuckovic et al., 2008). Moreover, the auricular branch of the vagus nerve projects afferently to the NTS which modulates heart rate only indirectly via the nucleus ambiguous and the dorsal vagal nucleus (Murray et al., 2016). Alternatively, taVNS indeed may not activate the vagus nerve in all cases, perhaps due to differences in stimulation amplitude, which would attenuate group-level effects compared to sham stimulation. However, brain imaging studies have consistently shown that taVNS activates the primary projection target of the vagal afferents, the NTS (Butt et al., 2020; Frangos et al., 2015; Yakunina et al., 2017), pointing to sufficiently robust effects at the group level. Another possibility is the lack of studies administering taVNS on the right side (four studies, three bilaterally so far), as cardio-vascular effects have been initially reported after right-sided stimulation (Burger et al., 2020). More specifically, the sinoatrial node is predominantly innervated by the right vagus (Ardell & Randall, 1986), and the only study stimulating the right side separately found increases in SDNN only for right-, but not for left-sided taVNS (De Couck et al., 2017). Although it is conceivable that more studies stimulating the right vagus nerve might change the results, vagal nerve endings in the ear are almost exclusively afferent (Kaniusas et al., 2019). Therefore information from both sides is mostly integrated centrally in the brainstem, before influencing HRV efferently (Chen et al., 2015). Nonetheless, more studies stimulating the right ear are necessary to conclusively resolve this question in the future. Regarding left-sided stimulation however, we have provided strong evidence supporting an absence of taVNS effects on HRV. Another open question is whether a different stimulation site than the auricle might affect HRV. Next to taVNS, transcutaneous vagus nerve stimulation can be applied cervically as well (tcVNS; transcutaneous cervical vagus nerve stimulation), i.e. in the neck adjacent to the carotid artery and target the cervical branch of the vagus nerve (Frangos & Komisaruk, 2017). As it is unclear whether tcVNS and taVNS produce similar physiological effects (Farmer et al., 2020), tcVNS might be able to increase HRV more consistently, as demonstrated in a study by Brock and colleagues (Brock et al., 2017). However, the number of studies reporting HRV changes in tcVNS is even smaller and more research is needed to conclusively address this possibility. To conclude, to identify potential biomarkers of taVNS, further research should focus on systemically investigating other neurophysiological signals, stimulation sides and electrode placements.

To synthesize as many studies as possible, we aggregated over different dependent measures of HRV. The resulting heterogeneity in conceptually related HRV measures might decrease the comparability of results, although the homogeneity of reported effect sizes speaks against this interpretation. Further, some of the measures have been questioned in their roles as useful proxies of HRV (Burger et al., 2020). For example, pNN50 has been criticized because of the arbitrariness of the 50ms cut-off (Mietus, 2002). Critically, SDNN reflects both sympathetic and parasympathetic influences and depends on the length of the recording period, thus making it difficult to compare SDNN measures obtained from recordings of different durations (Electrophysiology, 1996). Similarly, the LF/HF ratio reflects a mix of sympathetic and vagal activity and has a disputed physiological origin (Laborde et al., 2017) and thus might not clearly reflect HRV. Even though we excluded both SDNN and LF/HF ratio in the present analysis, they are included in the app as many authors still report them as proxies for HRV. However, we do not recommend including them in the interactive meta-analysis. Adopting standardized reporting protocols and identifying a best practice regarding taVNS research and HRV measurement is necessary (Borges et al., 2019; Farmer et al., 2020; Quintana et al., 2016). Still, a lack of standardized procedures characterizes many emerging fields. Therefore, our framework of a living Bayesian meta-analysis may help to quickly synthesize results to provide evidence-based guidance.

If HRV is not a suitable biomarker of acute taVNS, what might be a good alternative? Next to HRV, Burger and colleagues proposed other potential biomarkers, such as somatosensory evoked potentials (Fallgatter et al., 2003), pupil dilation, the P300 event-related potential, and salivary alpha-amylase (Warren et al., 2019). These candidates are predominantly based on the hypothesized effect of taVNS on the noradrenergic system. However, there is currently no conclusive evidence for their suitability as biomarkers with varying empirical success which could be due to the low power of many studies in the field (Burger et al., 2020). Another candidate is the electrogastrogram, which is supported by two studies so far (Teckentrup et al., 2020). Unfortunately, as the number of studies for alternative biomarkers is even lower than for HRV, a meta-analytic analysis is unlikely to yield conclusive results within the next few years. Recently, in-vivo recordings of the human vagus nerve have been demonstrated for the first time (Ottaviani et al., 2020) and this technique has immense potential to improve future taVNS research.

Despite its transparent and versatile format, this meta-analysis has limitations. First, taVNS serves as a potential treatment for patient groups where sympathetic dominance is part of the pathology, such as heart failure or atrial fibrillation (Akdemir & Benditt, 2016; Kaniusas et al., 2019). However, assuming generalizability of the results to clinical populations is problematic because of potential differences in baseline indices of cardiac vagal activity (Akdemir & Benditt, 2016). Second, except for a few studies (De Couck et al., 2017; Sclocco et al., 2020), earlobe sham is chosen as control condition, because it evokes a similar sensation to the participant, without actively targeting the vagus nerve (Frangos et al., 2015). Nonetheless, Butt and colleagues argue that the earlobe might not be the best sham stimulation site as it is not physiologically inert and produced similar patterns of activation to taVNS stimulation (Butt et al., 2020). However, as the earlobe is not innervated by the vagus nerve and earlobe stimulation evokes a similar sensation for participants, it provides a suitable sham stimulation site that may control for many potential confounds such as unblinding.

To conclude, emerging fields pose unique challenges in synthesizing evidence to guide future research efforts. By developing a living Bayesian meta-analysis, we can provide a comprehensive statistical framework to address the question whether HRV is a robust biomarker of taVNS based on the currently available evidence. Our results provide conclusive support for the null hypothesis that taVNS has no acute effect on HRV across a wide range of settings. Still, it is possible that taVNS could elicit effects in more restricted cases, for example, after stimulation of the right side or in patient populations with differences in baseline vagal activity. Crucially, our Shiny app provides a principled way to update the results as new studies will become available for an analysis. Such an open format is important in young research fields such as non-invasive brain stimulation with a limited number of studies that may still point towards promising clinical applications or help identify dead ends in research quickly. Ultimately, our approach allowed us to systematically aggregate evidence leading to conclusive evidence about a relevant research question that narrative reviews were unable to resolve. Therefore, future research may build on this worked example to formally synthesize the evidence to help settle debates in the field more effectively.

## Acknowledgement

VT & NBK received salary support from the University of Tübingen, fortune grant #2453-0-0. NBK received support from the Daimler and Benz Foundation, grant 32-04/19.

## Author contributions

NBK was responsible for the study concept and design. VW performed the literature search and AK & VW coded eligible studies. VW performed the data analysis and AK, VT, & NBK contributed to analyses. VW coded the corresponding shiny App. VW, AK, & NBK wrote the manuscript. All authors contributed to the interpretation of findings, provided critical revision of the manuscript for important intellectual content and approved the final version for publication.

## Footnotes

1 https://github.com/neuromadlab/taVNSHRVmeta

